## Supplemental Materials for "Functional specialization within the inferior parietal lobes across cognitive domains"

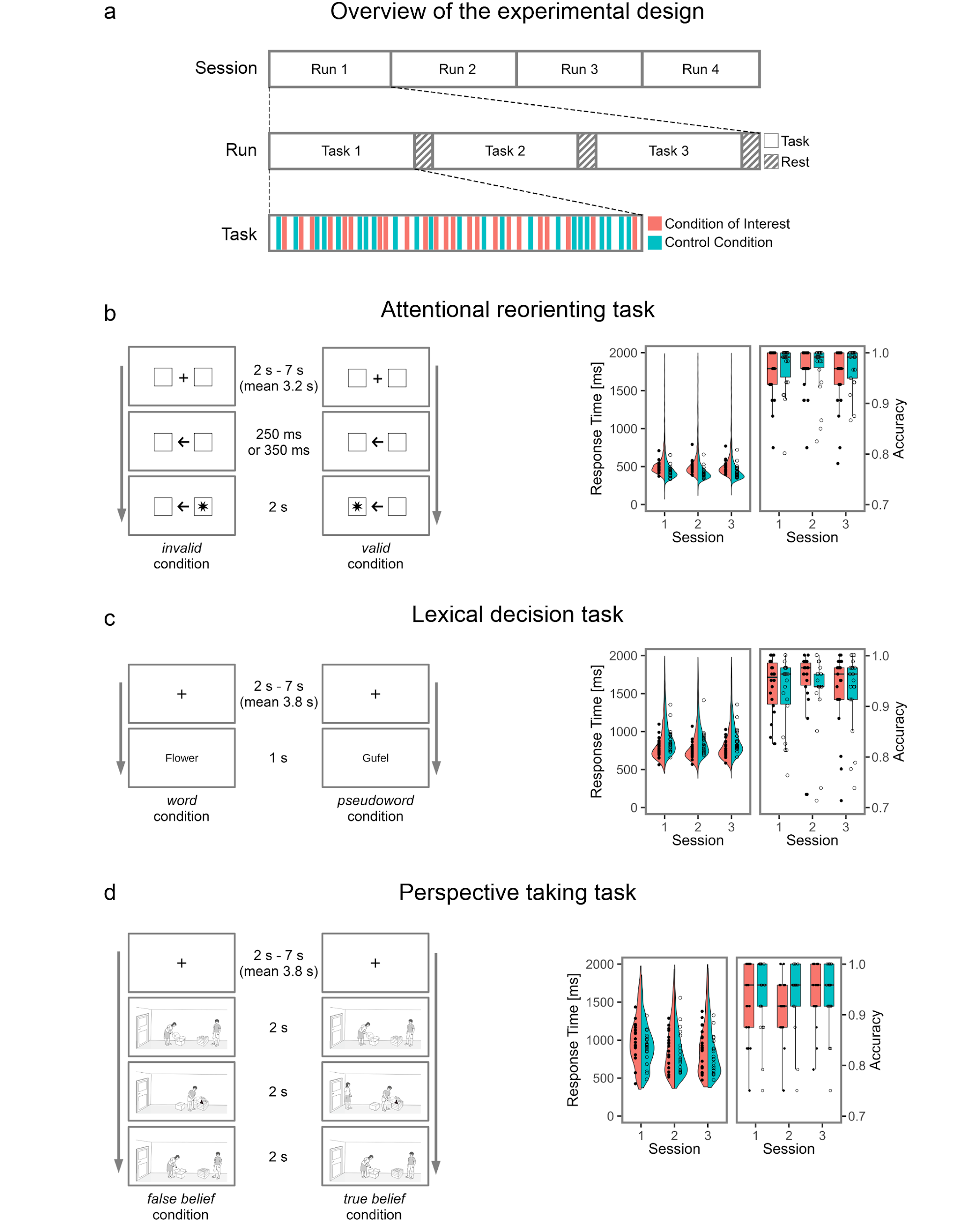

**Figure 1–figure supplement 1. Experimental design and behavioral results.** (**a) Experimental design.** Each of the four sessions consisted of four fMRI runs. In each run, all three tasks were presented in a pseudo-randomized order and analyzed in an event-related fashion. (**b) Attentional reorienting task.** In each trial, a directional arrow appeared at the center of the screen to direct the subject’s attention to the left or right. In 75 % of the trials, the arrow correctly predicted the position of the target (*valid* condition), in 20 % of the trials, the target appeared on the opposite side and subjects had to reorient their attention (*invalid* condition). In 5% of the trials, no response was prompted (*catch* condition). Subjects indicated the side of the target via button press. (**c) Lexical decision task**. Participants performed lexical decisions (*word* or *pseudoword* condition) based on concrete German nouns or well matched pseudowords. (**d) Perspective taking task.** In each trial, character A places a target object in a container. Thereafter, Character B changes the location of the target object. Character A has left the room (*false belief* condition) or watches the relocation of the object (*true belief* condition). Character A then searches for the target object at the location congruent with her / his knowledge (*expected*) or at the contradicting location (*unexpected*). Participants had to indicate via button press whether character A searched at the expected location or not. **Behavioral results** for each task are shown in the right panel, separately for each session. Single subject data is overlaid as circles.

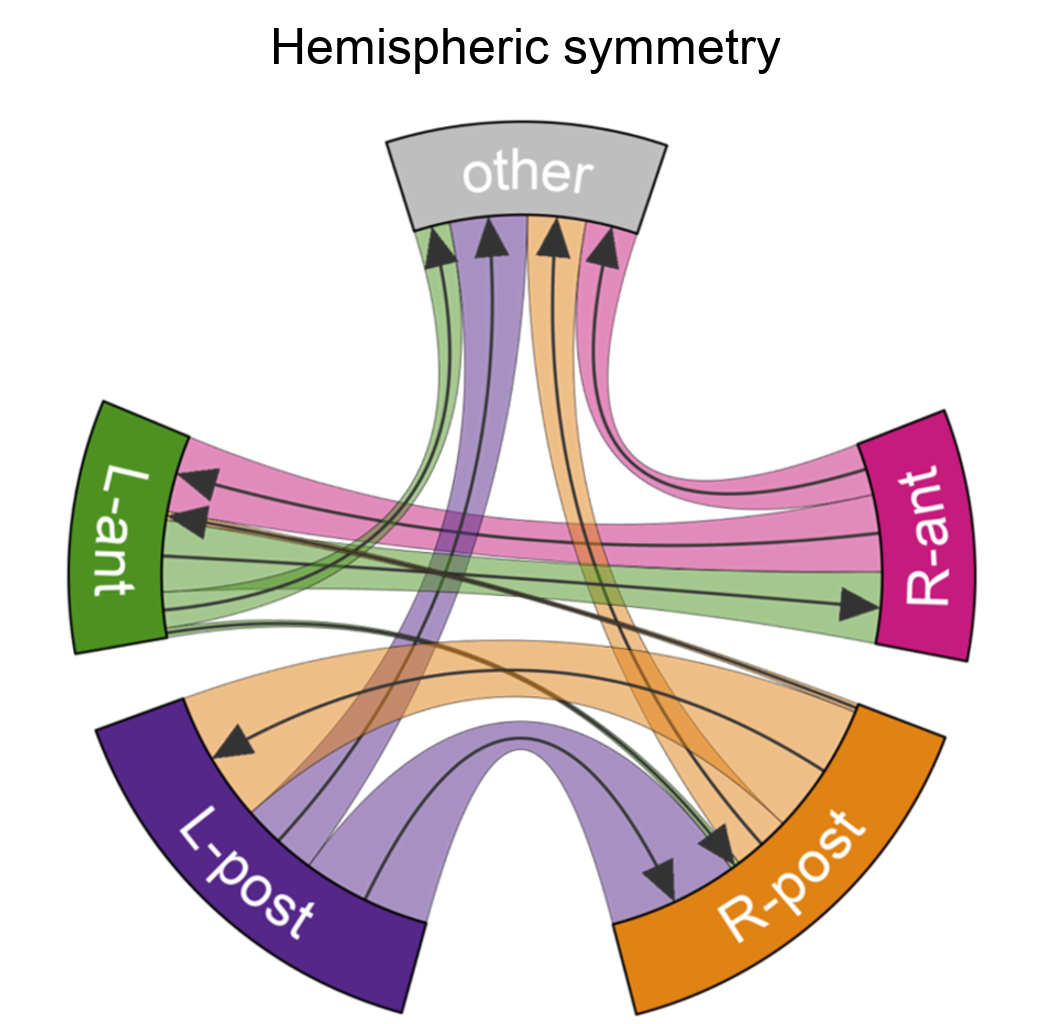

**Figure 2–figure supplement 1. The final subregion solution shows a high degree of symmetry between hemispheres.** Evidence supporting the hemispheric symmetry of the final two-subregion solution in the region of interest parcellation by clustering algorithms. Depicts which proportion of subregion-voxels falls into its homologue by flipping the x-axis, and which proportion falls into other areas.

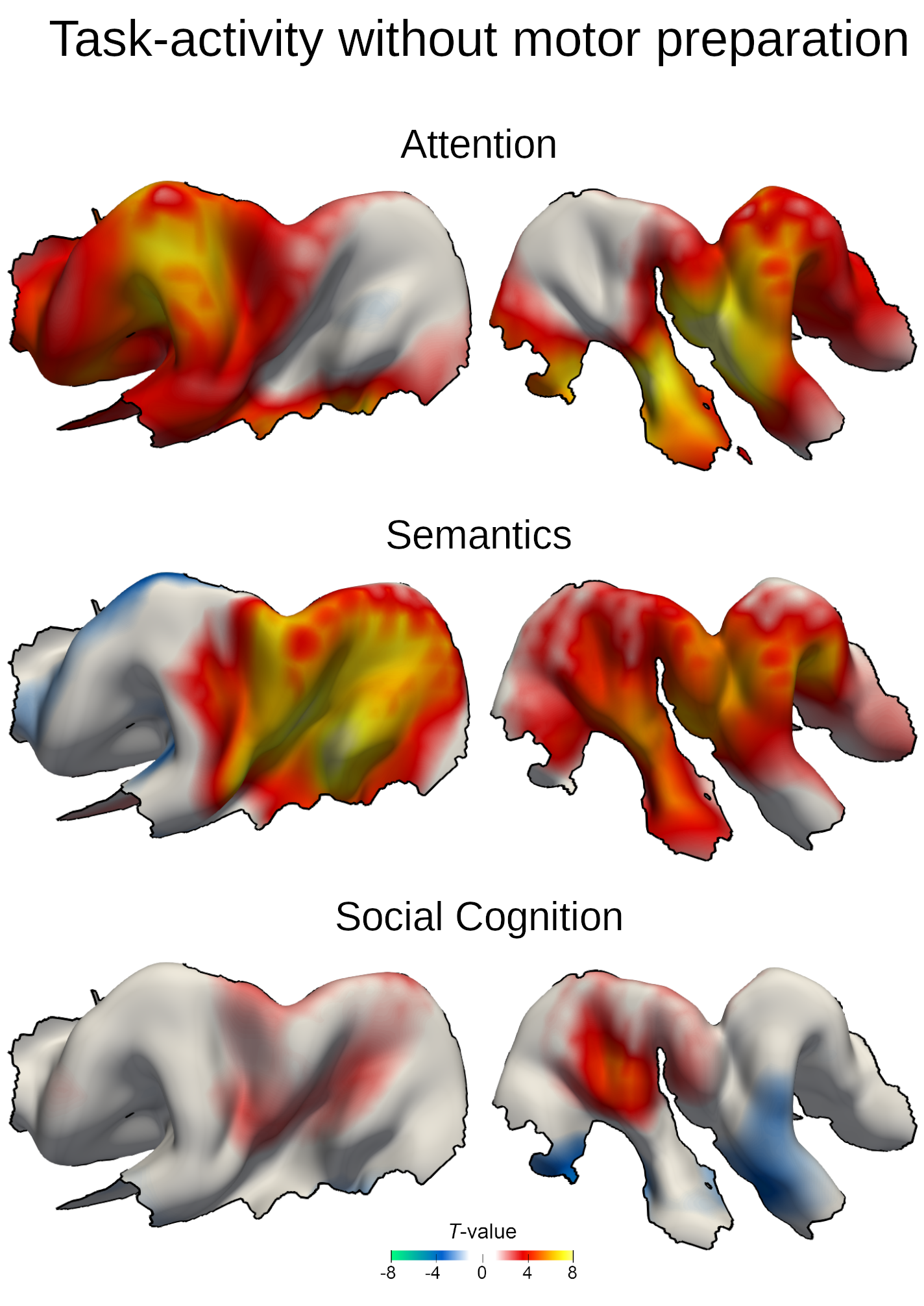

**Figure 2–figure supplement 2. Task-evoked neural responses in the IPL with explicit motor preparation modelling.** Task-dependent BOLD responses from the model that removes general motor preparation across tasks (GLM_cond+RT_) resemble results of the basic model (GLM_cond_, c.f. Figure 1). Colors indicate unthresholded T-values. Warm colors: higher GLM beta estimates for the target conditions. Cold colors: higher GLM beta estimates for the control condition.

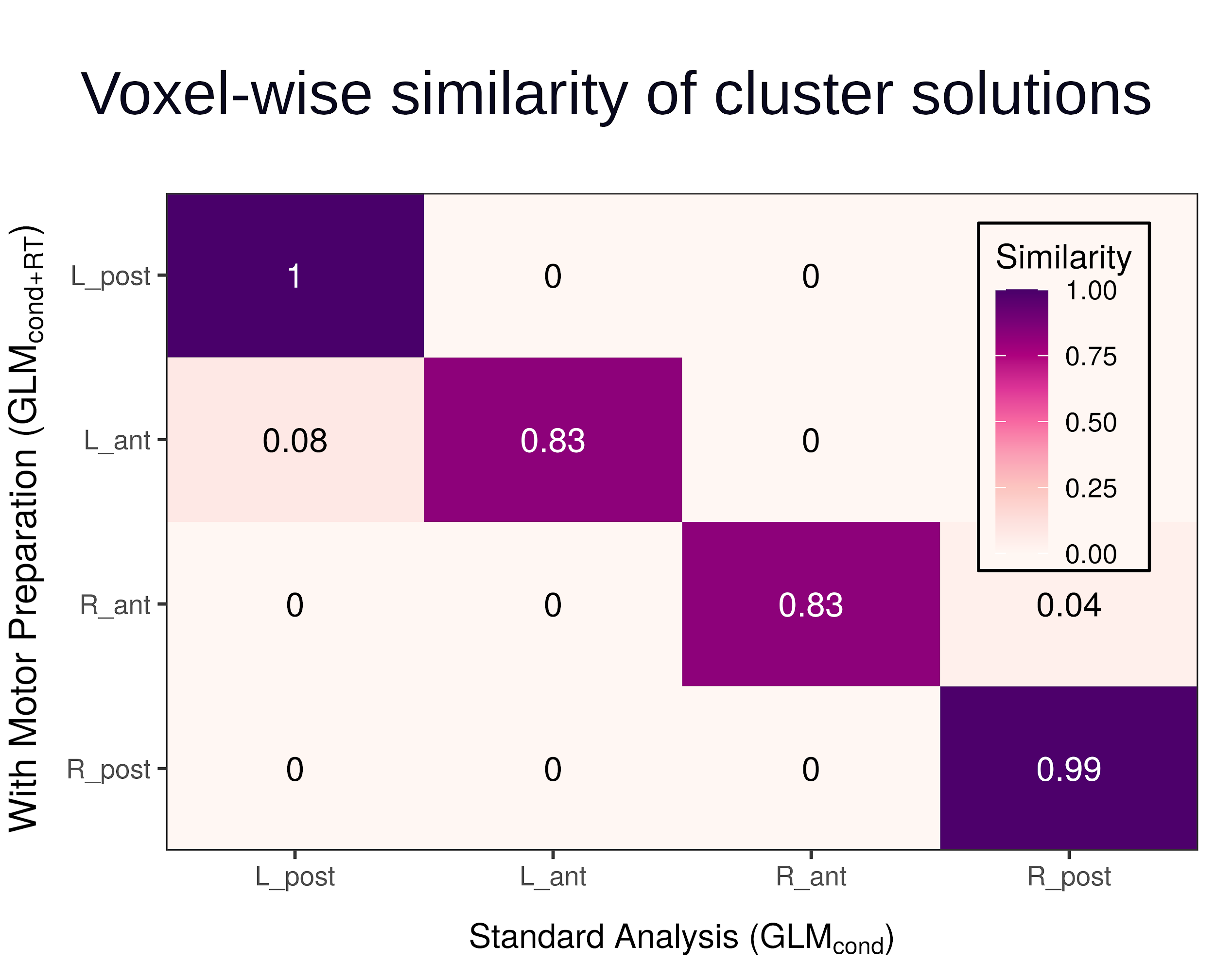

**Figure 2–figure supplement 3. Explicitly modelling motor preparation yields a similar clustering solution.** Depicts voxel-wise similarity of clusters across the final parcellation solutions between the basic model (GLM_cond_) and the model that removes general motor preparation across tasks (GLM_cond+RT_).

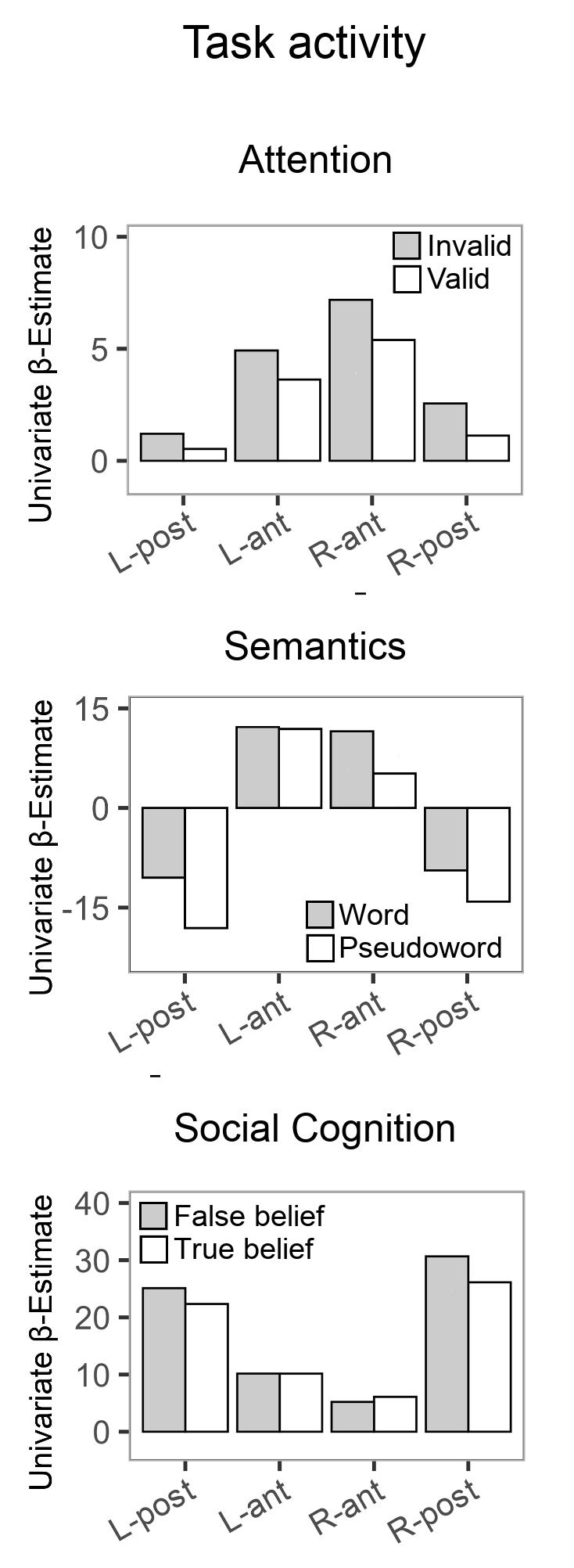

**Figure 3–figure supplement 1. Neural activity estimates for the target and control conditions of the three tasks.** For visualization purposes, beta estimates were extracted from GLM_cond_ at the center of mass for each IPL subregion.

**Figure 4–figure supplement 1. Functional connectivity between IPL subregions and large-scale brain networks.** The preferred connectivity between IPL subregions and brain-wide cortical regions identified task-specific coupling motifs of the four subregions. Large-scale brain networks were split into their left and right hemispheric parts. Functional connectivity was normalized per subregion and task. Anterior IPL subregions (top row) connect in a hemisphere specific way, while posterior IPL subregions (bottom row) connect in a more bilateral-symmetric way. LH/RH: left/right hemispheric network parts. See main text for details on the connectivity metric. Networks acronyms as defined in the main text.
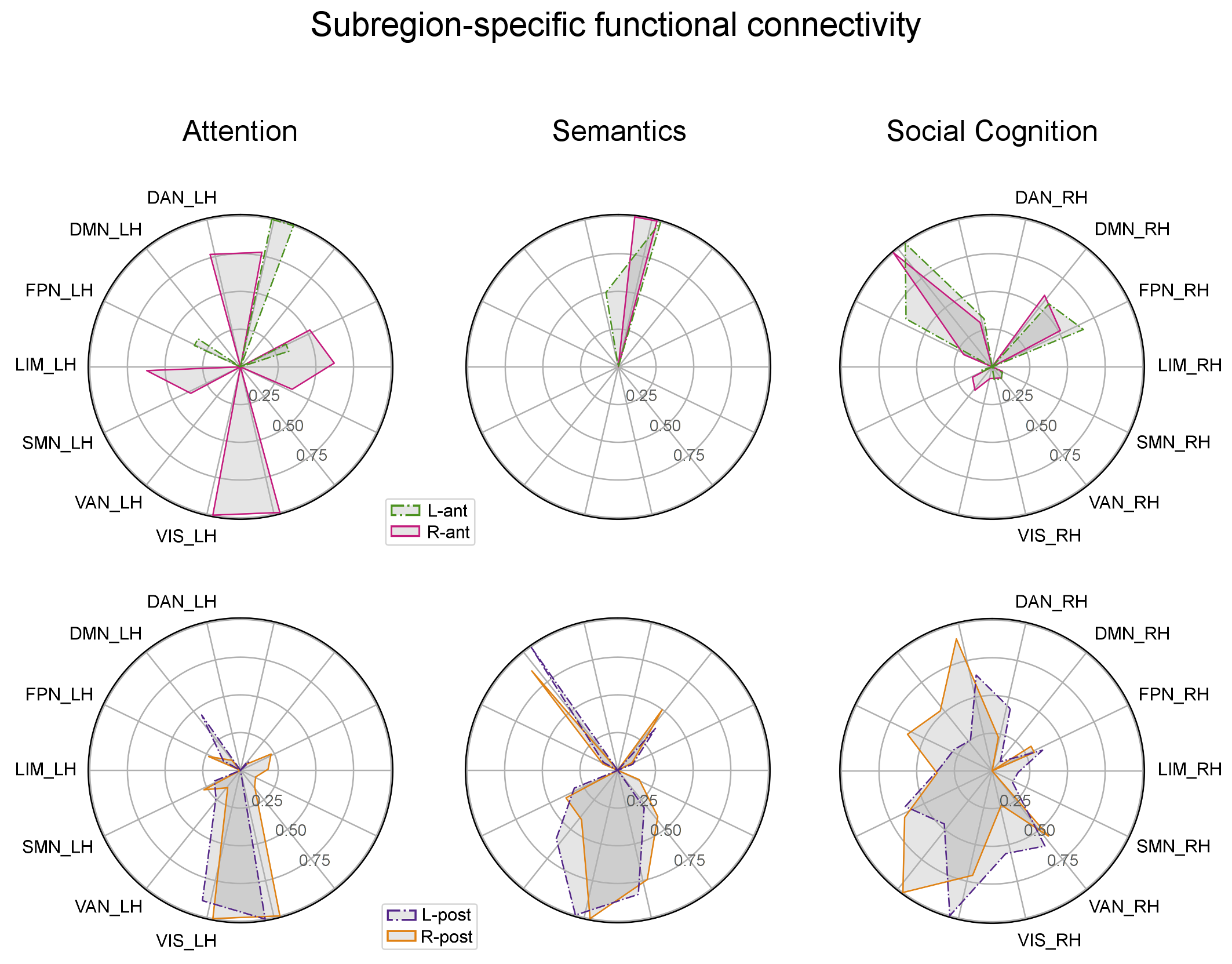

**Table S1**: Mass-univariate activations peaks for the three functional domains.

| **Region** | | **Anatomical assignment** | | **Hemis- phere** | **MNI coordinates  X Y Z** | | | **T-value** | **Cluster size** |
| --- | --- | --- | --- | --- | --- | --- | --- | --- | --- |
| Attentional reorienting | |  |  | |  |  |  |  |  |
|  | Supramarginal gyrus | PFm | R | | 57 | -48 | 24 | 9.68 | 181 |
|  | Superior temporal gyrus /   angular gyrus | PGa | R | | 63 | -45 | 17 | 9.23 |  |
|  | Middle temporal gyrus /   angular gyrus | PGp | R | | 48 | -63 | 14 | 8.42 |  |
|  | Middle temporal gyrus |  | R | | 57 | -57 | -1 | 7.03 |  |
|  | Precuneus | 7P | L | | 3 | -54 | 52 | 9.46 | 123 |
|  | Precuneus | 5L | R | | 6 | -60 | 59 | 8.27 |  |
|  | Precuneus | 7A | L | | 12 | -66 | 52 | 7.21 |  |
|  | Precentral gyrus |  | L | | -27 | -3 | 59 | 9.32 | 54 |
|  | Superior frontal gyrus |  | R | | 30 | -6 | 63 | 8.36 | 34 |
|  | Middle frontal gyrus |  | R | | 36 | -3 | 56 | 7.33 |  |
|  | Precentral gyrus |  | R | | 36 | 3 | 49 | 7.00 |  |
|  | Inferior parietal lobe | PF | L | | -54 | -42 | 38 | 7.35 | 33 |
|  | Postcentral gyrus |  | L | | -45 | -33 | 49 | 7.04 |  |
|  | Inferior parietal lobule | hIP2 | L | | -45 | -39 | 42 | 6.99 |  |
|  | Superior parietal lobe | hIP1 | R | | 30 | -48 | 42 | 10.10 | 31 |
| Lexical decisions | |  |  | |  |  |  |  |  |
|  | Angular gyrus | PGp | L | | -51 | -72 | 28 | 8.00 | 73 |
|  | Angular gyrus | PGp | L | | -39 | -75 | 42 | 7.36 |  |
|  | Angular gyrus | PGa | L | | -45 | -57 | 35 | 7.22 |  |
|  | Middle cingulate gyrus | 5M | L | | -6 | -30 | 42 | 9.07 | 50 |
|  | Superior frontal gyrus |  | L | | -18 | 36 | 49 | 7.94 | 34 |
|  | Middle frontal gyrus |  | L | | -27 | 24 | 52 | 6.64 |  |
| Perspective taking | |  |  | |  |  |  |  |  |
|  | Supplementary motor cortex |  | L | | -3 | 15 | 52 | 5.15 | 184 |
|  | Superior medial gyrus |  | L | | -6 | 24 | 42 | 4.37 |  |
|  | Superior frontal gyrus |  | L | | -15 | 12 | 63 | 3.84 |  |
|  | Posterior medial frontal g. |  | R | | 12 | 6 | 52 | 3.73 |  |
|  | Inferior frontal gyrus | BA45 | L | | -45 | 24 | 31 | 5.11 | 59 |
|  | Precentral gyrus |  | L | | -39 | 3 | 42 | 4.77 | 45 |
|  | not assigned |  | L | | -45 | 6 | 56 | 3.84 |  |
|  | Precentral gyrus | BA44 | L | | -48 | 9 | 35 | 3.81 |  |
|  | Middle frontal gyrus |  | L | | -42 | 3 | 52 | 3.72 |  |
|  | Angular gyrus | PGa | R | | 51 | -60 | 31 | 5.15 | 32 |
|  | Precuneus |  | R | |  |  |  | 4.50 | 31 |
|  | Precuneus |  | L | |  |  |  | 3.98 |  |
|  | Middle temporal gyrus |  | R | | 54 | -18 | -15 | 4.68 | 20 |
|  | Middle temporal gyrus |  | R | | 60 | -15 | -11 | 4.29 |  |

*Note: Attention and semantic tasks: thresholded at p=0.05, FWE corrected, cluster extent ≥20 voxels. Social cognition task: thresholded at p=0.001, uncorrected, cluster extent ≥20 voxels. Anatomical assignment according to SPM Anatomy toolbox (v. 22c).*

**Table S2**: Intrinsic effective connectivity and task-specific self-connectivity modulation

| **Source subregion** |  | **Target subregion** | **Strength** | **95%-CI** |
| --- | --- | --- | --- | --- |
| **Intrinsic connectivity** | | | | |
| L-ant | → | L-ant | -1.20 | [-1.21; -1.20] |
| L-ant | → | L-post | -0.37 | [-0.38; -0.37] |
| L-ant | → | R-ant | -0.33 | [-0.33; -0.33] |
| L-ant | → | R-post | -0.56 | [-0.56; -0.56] |
| L-post | → | L-ant | -0.05 | [-0.06; -0.05] |
| L-post | → | L-post | -0.79 | [-0.81; -0.79] |
| L-post | → | R-ant | 0.15 | [0.15; 0.16] |
| L-post | → | R-post | -0.29 | [-0.29; -0.29] |
| R-ant | → | L-ant | -0.27 | [-0.27; -0.27] |
| R-ant | → | L-post | 0.38 | [0.39; 0.39] |
| R-ant | → | R-ant | -1.13 | [-1.14; -1.13] |
| R-ant | → | R-post | 0.87 | [0.87; 0.88] |
| R-post | → | L-ant | 0.34 | [0.34; 0.35] |
| R-post | → | L-post | 0.42 | [0.42; 0.43] |
| R-post | → | R-ant | -0.30 | [-0.30; -0.30] |
| R-post | → | R-post | -1.14 | [-1.15; -1.14] |
| **Source subregion** |  | **Target subregion** | **Modulation strength** | **p-value** |
| **Self connectivity** | | | | |
| Semantics | | | | |
| L-ant | → | L-ant | -5.53 | 0.0006 |
| L-post | → | L-post | -6.88 | 0.0010 |
| Social Cognition | | | | |
| R-ant | → | R-ant | -4.01 | 0.0055 |

*Note: Intrinsic connectivity parameters (‘A-matrix’) and significant modulatory parameters (‘B-matrix’,* α ≤ 0.01*) of self-connections. Strength is given as posterior expectation from the optimum Bayesian parameter average model. The 95% confidence intervals were built from the corresponding Bayesian parameter covariances and exceed zero for all parameters. P-values for self-connectivity are based on a random effects permutation test for the null hypothesis ‘no parameter difference between tasks’.*
